# No Effect of Exercise Timing on Daily Glycaemia Despite Shifts in Acute Glucose Dynamics in Participants with Normal Glucose Tolerance: A Randomised Crossover Trial

**DOI:** 10.64898/2026.10.30.763114

**Authors:** Lara T. Dasar, Matthew Hamilton, Daria Borisova, Imogen Lewis, Victoria Kjellin-Bennett, Vanessa Gargano, Ratnika Prakash, Kristine Beaulieu, Brendan M. Gabriel

**Author notes:** Corresponding author: Brendan M. Gabriel – School of Medicine, Medical Science and Nutrition, Rowett Institute of Nutrition and Health, University of Aberdeen, Aberdeen, UK.

## Abstract

Exercise is a well-established modulator of glucose regulation, but whether the time of day it is performed matters for healthy, free-living adults remains unresolved. In a randomised crossover trial, 29 healthy adults completed a single 30-minute bout of moderate-intensity exercise in the morning (07:00–10:00) and in the evening (16:00–19:00), with continuous glucose monitoring and wrist-worn activity tracking used to capture responses over an 8-day free-living protocol. Evening exercise produced a significantly greater acute reduction in blood glucose (*P* < 0.001) compared to morning exercise, but this did not translate into consistent differences in overall daily mean glycaemic exposure, whether measured over the calendar day or over a window aligned with exercise onset. Evening exercise was linked to greater within-day glycaemic variability than morning exercise, with the clearest evidence coming from the circadian-aligned analysis (day × timing interaction, SD: *P* = 0.00939; CV: *P* = 0.00766). Appetite rose after exercise in both conditions, with no difference by time of day. Taken together, these results show that exercise timing does not change how much glucose exposure a healthy adult accumulates over a day, but it does change the shape of the glucose response around the exercise bout itself, a distinction only visible when glucose is examined at a finer temporal resolution than daily averages allow.

## 1 Introduction

Metabolic health is shaped by interactions between behaviour, physiological regulation, and internal timing systems. Exercise is a well-established modulator of glucose homeostasis and appetite, yet accumulating evidence suggests that the time of day at which exercise is performed may influence its metabolic effects (Schroder et al., 2015; Smith et al., 2025).

This interest in *chrono-exercise* is supported by circadian variation in skeletal muscle oxidative capacity, insulin sensitivity, substrate utilisation, and appetite-related signalling, all of which fluctuate across the day (Gabriel et al., 2021; Templeman et al., 2021; Van Moorsel et al., 2016). Given the tight coupling between glycaemic regulation and appetite control, understanding how exercise timing affects both systems may provide a more integrated view of metabolic responses across the day.

Circadian disruption is a common feature of metabolic dysfunction whereby the intrinsic rhythm of skeletal muscle insulin sensitivity is blunted, and time-of-day-targeted exercise interventions could help restore circadian alignment (Hesketh, 2025). Much of the existing chrono-exercise literature, therefore, has focused on individuals with obesity, insulin resistance, or impaired glycaemic control, where afternoon or evening exercise often leads to more favourable glycaemic responses compared with morning exercise (Mancilla et al., 2021; Moholdt et al., 2021; Qian et al., 2023; Savikj et al., 2019, 2022; Van Der Velde et al., 2023). These findings have generated interest in leveraging circadian physiology to enhance metabolic benefits where regulation is compromised.

The metabolic significance of exercise timing in healthy individuals appears to have received less attention. Characterising how exercise timing interacts with endogenous rhythms in normoglycaemic adults is important for understanding the fundamental physiology of chrono-exercise, informing general physical activity guidelines, and establishing a reference point against which clinical populations can be compared. Findings in this population are mixed, and most studies have been conducted under tightly controlled laboratory settings rather than free-living conditions (Kim et al., 2022; Rodriguez-Muñoz et al., 2024; Tanaka et al., 2021). Nonetheless, there is a growing need for timing to be considered as a key principle of exercise prescription alongside frequency, intensity, time, and type (Reid et al., 2019; Ruiz et al., 2024). With increasing access to wearable technologies, free-living research is an opportunity to characterise real-world responses to time-targeted exercise. For example, using continuous glucose monitors (CGMs) and physical activity monitors to capture dynamic glycaemic profiles and real-time activity may enhance exercise timing strategies based on real-world settings (Chang et al., 2023; Shandhi et al., 2021). Engagement with these technologies may reveal subtler physiological responses that would otherwise be obscured in laboratory settings, and may help address whether acute timing-related differences translate into meaningful long-term metabolic effects.

Against this background, this randomised crossover trial examined the effects of morning versus evening moderate-intensity exercise on acute and daily glycaemic patterns and subjective appetite in healthy adults living in their habitual environment. By capturing responses under ecologically valid free-living conditions, this study provides foundational evidence for understanding whether exercise timing influences glycaemic regulation and appetite control in healthy individuals, and offers insight into designing more personalised, time-sensitive exercise prescriptions.

## 2 Methods

### 2.1 Ethical Approval

Informed consent was received from all participants prior to taking part in the study. Study procedures were conducted in accordance with the *Declaration of Helsinki* and were approved by the Rowett Ethics Panel of The University of Aberdeen. The full protocol is available on the Open Science Framework (DOI: 10.17605/OSF.IO/6TB78).

### 2.2 Participants

Participants were recruited via distribution of posters and emails, and social media advertising. Eligible participants were healthy adults, aged ≥18 years with a BMI < 28 kg/m^2^. Screening sessions included anthropometric measurements (weight, height and blood pressure) and ensuring that participants were safe to undergo exercise, assessed using a Physical Activity Readiness Questionnaire (PAR-Q) (Humphrey & Lakomy, 2003). Potential participants were excluded if they met any of the following criteria: claustrophobia, nicotine use (current or within the past 6 months), cardiovascular condition, rheumatic illness, cancer, psychiatric disorder, metabolic disease, unable to speak or understand English, or blood pressure consistently <90/60 mmHg or >140/90 mmHg.

Sample size estimation was based on glycaemia using effect size estimates from a comparable crossover trial (Peña Carrillo et al., 2024). Using a mean difference of 0.69 mmol·L ¹, SD 1.32 mmol·L ¹, α = 0.05 and a paired design, 27 participants provide approximately 80% power to detect moderate differences in daily glucose outcomes between morning and evening exercise.

Participants were block randomised to start with morning exercise or evening exercise, before completing the other on days 4 and 8.

### 2.3 Experimental Protocol

Before the 8-day study, participants completed the Morningness-Eveningness Questionnaire (MEQ) to determine chronotype (Horne & Östberg, 1976). Participants completed the 19-item questionnaire, with each item’s score converted to a global score on a 5-point scale: definitely morning type (70–86), moderately morning type (59–69), neither type (42–58), moderately evening type (31–41), and definitely evening type (16–30). Participants were asked to refrain from other strenuous exercise (e.g. running, weight training, and sports) during the 8 days and asked not to consume alcohol during this time, as well as 24 hours prior to the study start date.

Participants completed two 30-minute exercise sessions in a randomised crossover fashion, one in the morning (between 07:00–10:00), and one in the evening (between 16:00–19:00), on day 4 and day 8, with days 1-3 and 5-8 (washout) as their normal routine. Both exercise sessions were conducted as follows. Participants were asked not to consume food or caffeine during the two-hour period before, or during exercise. Exercise was performed on a self-selected outdoor route which participants were advised to replicate across both sessions, comprising a mix of walking and running. To meet a moderate exercise intensity, participants were provided with a target heart beats per minute (bpm), calculated as 70% of age-predicted HRmax (70% of age-predicted HRmax (0.70 × [220 − age]) (American College of Sports Medicine, 2014). Participants were asked to match this target by monitoring HR on a wrist-worn physical activity monitor (Garmin Vívosmart 5, Garmin Ltd., Kansas, USA) which was worn continuously throughout the study, recording other parameters such as activity levels.

Immediately pre- and post-exercise, appetite was assessed using 100-mm Visual Analogue Scales (VAS) (Blundell & Rogers, 1980; Hill & Blundell, 1982), a validated measure of motivation to eat (Dakin et al., 2026). Participants rated subjective appetite sensations such as hunger, fullness, desire to eat and prospective food consumption to produce a composite appetite score (CAS). CAS was calculated from a standardised formula (Zafar et al., 2011) with higher scores indicating greater subjective appetite: CAS = [hunger + desire to eat + prospective food consumption + (100 – fullness)]/4.

Throughout the 8 days, participants wore a continuous glucose monitor (CGM; FreeStyle Libre 2, Abbott, Witney, UK) to measure blood glucose and completed daily estimated food diaries. Female participants answered menstrual-related questions on the Garmin Connect phone application (Appendix Table A1). Figure 1 shows a schematic diagram of the study timeline.

**FIGURE 1:**
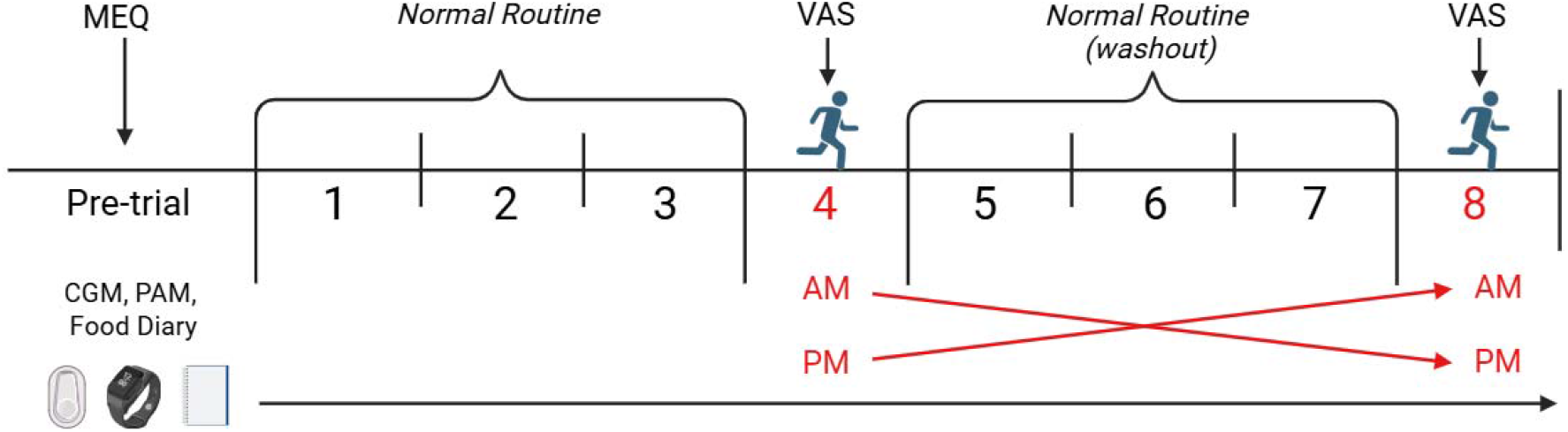
Summary of the 8-day randomised crossover design. Participants exercised for 30 minutes at moderate intensity (70% age-predicted HRmax) on day 4 and day 8. Morning exercise occurred between 07:00– 10:00, and evening exercise between 16:00–19:00. PAM; physical activity monitor. Created in https://BioRender.com.

### 2.4 Data Analysis

All statistical analyses were performed using R Version 4.4.1 (*R Core Team*, 2024). Statistical significance was set at *P* ≤ 0.05. Descriptive data are reported as mean ± SD, unless otherwise stated. Data visualisation was conducted using the *ggplot2* package.

Adherence between exercise conditions was assessed for exercise intensity (HR) using paired analysis. The Wilcoxon signed-rank test was applied when violations of normality were observed using the Shapiro–Wilk test and, and the data are presented as median (IQR). Effect sizes were calculated as *r* and interpreted according to standard thresholds (small: 0.10, moderate: 0.30, large: 0.50; (Field, 2009).

Analyses of blood glucose and appetite between exercise conditions were performed using linear mixed models (LMMs) implemented with the packages *lme4* and *lmerTest*. Model assumptions were evaluated using residual diagnostics, including Q–Q plots and assessment of homoscedasticity. LMMs were fitted using restricted maximum likelihood and included participant as a random effect to account for repeated measures within individuals, except where an alternative random-effects structure was specified. Where likelihood ratio tests indicated that a random slopes model significantly improved fit and did not produce a singular fit, the structure was retained. Log transformation was applied where necessary to satisfy model assumptions, with back-transformed geometric mean estimated marginal means (EMMs) reported in these cases. Where model residuals significantly deviated from normality, and a log transformation was not applied, bootstrap resampling (1,000 replicates) was conducted to validate parametric conclusions. Overall main effects and interactions were assessed using Type III Analysis of Variance with Satterthwaite’s method. Where a significant interaction was identified, post hoc contrasts were conducted using the *emmeans* package to examine differences within each exercise timing condition separately. Confidence intervals (CIs) for post hoc contrasts were calculated using the Wald method based on the Kenward-Roger degrees of freedom. Results are reported as EMMs with 95% CIs.

Acute glucose (model 1) and CAS (model 2) were analysed using pre- and post-exercise measurements that were collected before and after exercise. Both models included exercise timing (morning *vs.* evening), timepoint (pre *vs.* post), and their interaction as fixed effects.

Daily blood glucose was analysed using two complementary analytical windows. The calendar day window (midnight to midnight) (model 3) captured overall daily glycaemic load irrespective of when exercise occurred within the day. The circadian-aligned window (model 4) was anchored to the time of exercise onset, enabling assessment of glucose dynamics relative to the physiological stimulus of exercise. Baseline for both approaches was defined as the 24-hour window on the day preceding exercise. Both models included exercise timing (morning *vs.* evening), day (baseline *vs.* exercise day), and their interaction as fixed effects. Mean glucose, glucose SD, and glucose CV were calculated for each participant-day combination across both approaches. CGM coverage was estimated assuming 96 readings per 24 hours, consistent with the 15-minute sampling interval of the devices used. All participant-days were retained in primary analyses regardless of coverage, with a ≥70% coverage threshold examined in sensitivity analyses.

Sensitivity analyses were conducted for all primary models to assess the robustness of the findings. Feeding state (fed *vs.* fasted) was included as a covariate in sensitivity analyses for models 1 and 2, and mean glucose in models 3 and 4. SD and CV reflect within-day variability rather than absolute glucose concentration and therefore, feeding state was not included as a covariate for these outcomes in models 3 and 4. Additional sensitivity analyses incorporating sex, age, BMI, season, and chronotype were conducted for models 1 and 2. As none of these covariates significantly predicted acute glucose or appetite, they were not extended to models 3 and 4.

Participants were excluded if CGM data indicated time above range (TAR; >10.0 mmol·L ¹) based on a target of <25% for individuals with type 1 and type 2 diabetes (Battelino et al., 2019). A more stringent threshold of 5% was applied in the present study given that participants were healthy adults, consistent with normative CGM data indicating that this population typically spends approximately 1.2% of time above this level (Spartano et al., 2025).

## 3 Results

### 3.1 Participants

Of the 53 participants assessed for eligibility, 19 did not meet the inclusion criteria. Four participants withdrew after being randomised and prior to study onset due to time commitment and personal health issues. A total of 30 participants completed the study, 12 (9 female, 3 male) of which were allocated to group A (morning/evening) and 18 (10 female, 8 male) to group B (evening/morning). Participants who commenced exercise more than 30 mins after the prescribed time window were excluded from analysis (*n* = 1: female), a threshold selected based on the approximate hourly timescale of circadian variation in metabolic parameters, such that deviations beyond this point could meaningfully alter the circadian context.

The final analysis included 29 healthy adults (11 male and 18 female). Two male participants were excluded from model 2 due to data loss in CAS across all conditions, and one male participant was excluded from sensitivity analyses in models 3 and 4 due to <70% CGM coverage. Figure 2 shows a *Consolidated Standards Of Reporting Trials* (CONSORT) diagram of the recruitment process. Baseline participant characteristics are included in Table 1.

**FIGURE 2:**
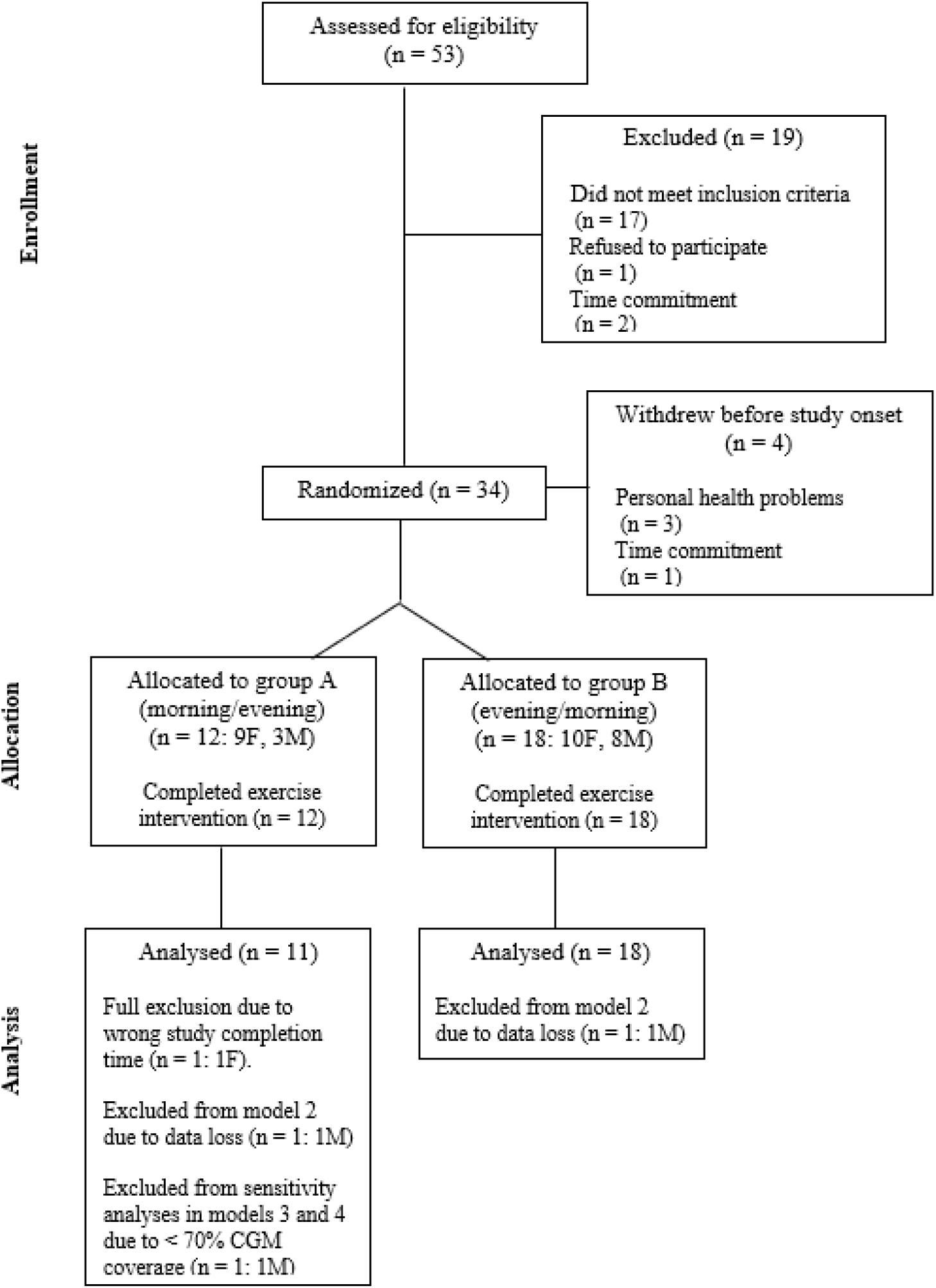
CONSORT diagram of recruitment, randomisation and analysis. Group A = morning then evening exercise; group B = evening then morning exercise. F = female; M = male. Created in https://BioRender.com.

**TABLE 1.** Participant Characteristics.

|  | **Female (*n* = 18)** | **Male (*n* = 11)** | **Total (*n* = 29)** |
|---|---|---|---|
| **Age (years)** | 35 ± 13 | 32 ± 11 | 35 ± 12 |
| **Body Mass (kg)** | 63.4 ± 8.1 | 73.3 ± 10.8 | 67.4 ± 10.3 |
| **Height (m)** | 1.65 ± 0.06 | 1.78 ± 0.07 | 1.70 ± 0.09 |
| **BMI (kg/m²)** | 23.2 ± 2.2 | 23.2 ± 2.4 | 23.0 ± 2.3 |
| **Estimated 70% MHR (bpm)** | 130 ± 9 | 131 ± 8 | 132 ± 8 |
| **Chronotype n (%)**<sup>1</sup> |  |  |  |
| Definitely morning type | 2 (11.1) | 0 (0) | 2 (7.1) |
| Moderately Morning Type | 3 (16.6) | 4 (40) | 7 (25) |
| Neither Type | 12 (66.7) | 6 (60) | 18 (64.3) |
| Moderately Evening Type | 1 (5.6) | 0 (0) | 1 (3.6) |

Data represent mean ± SD unless otherwise stated.
<sup>1</sup> Chronotype data unavailable for one male participant (*n* = 28).

### 3.2 Free-Living Exercise and Adherence

All 29 participants completed both exercise sessions and engaged with the wearable technologies. The average start time of morning exercise was 08:39 and the average evening time was 17:32, indicating that participants performed exercise within the prescribed windows (07:00–10:00 and 16:00–19:00).

All 29 participants completed food diaries. Nine participants did not adhere to the pre-exercise eating protocol (no food or caffeine during the two-hour period before exercise), across 6 morning sessions (and hence not fasted) and 6 evening sessions. A further 2 participants did not complete morning exercise in a fasted state (i.e. they consumed food/caffeine 2 or more hours before the start of exercise and still adhered to the pre-exercise eating protocol). One participant’s last consumption was at 06:45 and the other was at 04:00. All 29 participants were included in a paired analysis. Subsequently, 21 participants completed morning exercise in a fasted state, with an average time of last intake the evening prior of 20:32. All participants completed evening exercise in a fed state. Visual inspection (histogram and Q-Q plot) and the Shapiro-Wilk test indicated that the differences in intake were normally distributed (W = 0.96, *P* = 0.3), so a paired t-test was used. Intake did not differ between morning exercise 2151 ± 1112 and evening exercise 2095 ± 902, *t*(28) = 0.32, *P* = 0.75, mean difference = 55.72 (-303.81, 415.26), *dz* = 0.059.

The physical activity monitor showed reliable activity tracking performance, successfully capturing HR from 51 out of the 58 exercise sessions (∼88%). Among participants with paired values, adherence to the prescribed exercise intensity (70% of age-predicted HRmax) was assessed, which compared mean %HRmax between morning and evening (*n* = 23).

The Shapiro–Wilk test indicated non-normality (W = 0.91, *P* = 0.0317) and therefore, a non-parametric test was applied. A Wilcoxon signed-rank test showed no significant difference between morning and evening sessions (*V* = 129, *P* = 0.948) and a negligible effect size (*r* = 0.01). Median % HRmax was 73.6% (70.5,77.6) in the morning condition and 72.9% (71.2, 79.4) in the evening (Figure 3).

**FIGURE 3:**
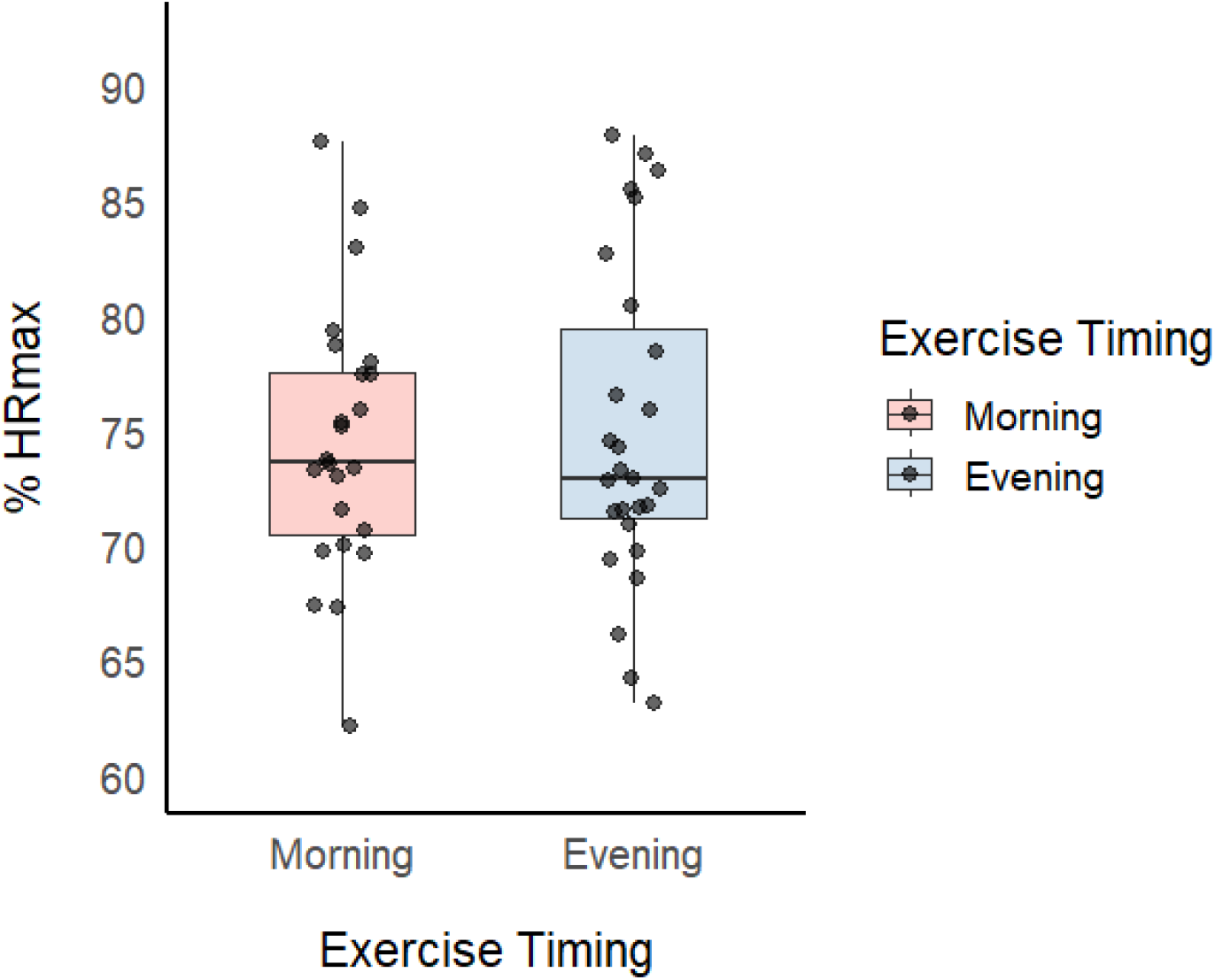
Exercise intensity between time-conditions (*n* = 23). Exercise intensity expressed as % HRmax. No statistical differences observed (*P* = 0.948) between morning and evening exercise conditions using a Wilcoxon signed-rank test. Boxplots represent median and IQR.

### 3.3 Model 1: Acute Blood Glucose

Blood glucose was analysed during the 30-minute period before (pre) and after (post) exercise. Data from 29 participants were included in a primary analysis, yielding 106 observations across four condition cells (morning/evening × pre/post). Five participants had two missing glucose observations, resulting in a total of 10 missing values, which was accommodated for by the mixed-effects framework. The LMM (Model 1, Figure 4) included exercise timing (morning *vs.* evening), timepoint (pre *vs.* post), and their interaction, with participant included as a random effect (*n* = 29).

**FIGURE 4:**
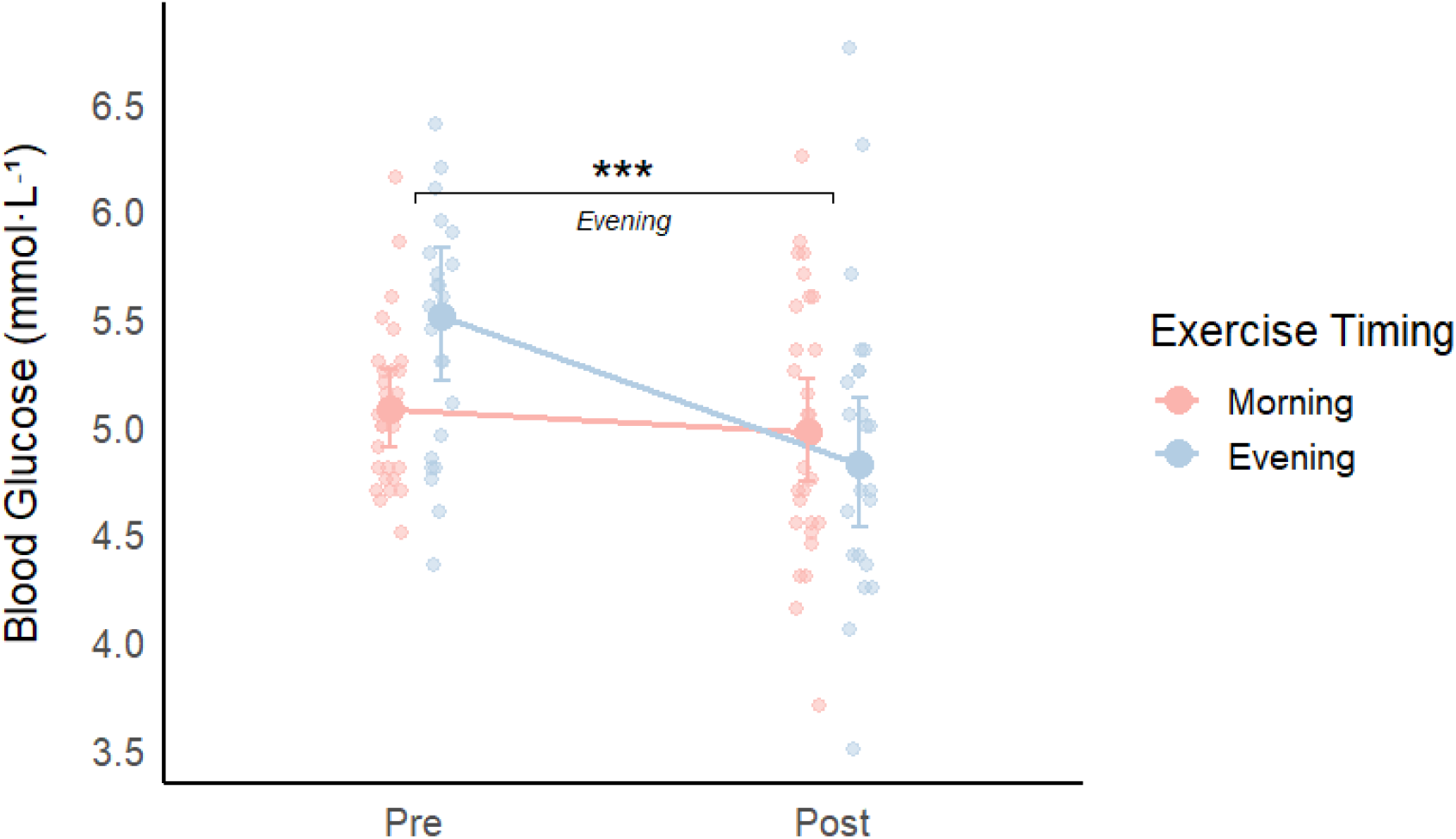
Blood glucose concentrations pre and post exercise in the morning and evening conditions (*n* = 29). There was a significant exercise timing × timepoint interaction (F(1, 25.344) = 13.5473, *P* = 0.001101). Blood glucose decreased significantly from pre- to post-exercise in the evening condition (pre: 5.51 mmol·L ¹, 95% CI [5.21, 5.83]; post: 4.82 mmol·L ¹, 95% CI [4.53, 5.13], *P* < 0.001), from pre- to post-exercise in the morning condition (pre: 5.07 mmol·L ¹, 95% CI [4.90, 5.26]; post: 4.98 mmol·L ¹, 95% CI [4.74, 5.22], *P* = 0.4452). *** *P* < 0.001, evening condition only. Data presented as back-transformed EMMs ± 95% CIs with individual data points.

Performing residual diagnostics indicated that there was a violation of normality (Shapiro– Wilk: W = 0.93, *P* < 0.001), driven by one elevated glucose observation (9.15 mmol·L ¹). Log transformation resolved this violation (Shapiro–Wilk: W = 0.98, *P* = 0.0681) and was therefore applied to subsequent analyses. A likelihood ratio test indicated that a random slopes model significantly improved model fit on the log-transformed outcome (χ²(5) = 13.05, *P* = 0.023), and the random slopes model did not produce a singular fit. Residual diagnostics confirmed that model assumptions were satisfied in the random slopes model (Shapiro–Wilk: W = 0.99, *P* = 0.519) and was therefore retained as the final model.

There was no significant main effect of exercise timing (F(1, 24.78) = 1.08, *P* = 0.309). The model revealed a significant main effect of timepoint (F(1, 26.27) = 12.86, *P* = 0.00135) and exercise timing × timepoint interaction (F(1, 25.34) = 13.55, *P* = 0.00110).

Fixed-effect estimates showed that pre-exercise glucose concentrations were higher in the evening compared to the morning condition (β = 0.08, SE = 0.029, *P* = 0.00681). The significant exercise timing × timepoint interaction (β = −0.11, SE = 0.03, *P* = 0.00110) confirmed that the reduction in glucose from pre- to post-exercise was greater in the evening condition relative to the morning condition. Back-transformed geometric mean EMMs indicated that in the evening, pre-exercise glucose was 5.51 mmol·L ¹ (95% CI [5.21, 5.83]) and post-exercise glucose was 4.82 mmol·L ¹ (95% CI [4.53, 5.13]). In the morning, pre-exercise glucose was 5.07 mmol·L ¹ (95% CI [4.90, 5.26]) and post-exercise glucose was 4.98 mmol·L ¹ (95% CI [4.74, 5.22]).

Post hoc analyses revealed that blood glucose decreased significantly from pre- to post-exercise in the evening condition (log-scale contrast: estimate = 0.134, 95% CI [0.079, 0.188], *P* < 0.001). No significant change was observed from pre- to post-exercise in the morning condition (estimate = 0.020, 95% CI [−0.032, 0.072], *P* = 0.445), indicating that the acute blood glucose response differed by timing.

Sensitivity analyses were performed following removal of one elevated glucose observation (9.15 mmol·L ¹). Residual diagnostics confirmed that assumptions were satisfied with a random slopes structure and log transformation (Shapiro–Wilk: W = 0.99, *P* = 0.467).

Removal of the observation did not alter the primary findings as the exercise timing × timepoint interaction remaining statistically significant (F(1, 24.114) = 8.785, *P* = 0.00674). The observation was therefore retained in final analyses.

A sensitivity model including feeding state (fed *vs.* fasted) as a covariate did not significantly improve model fit (χ²(1) = 0.03, *P* = 0.864), and feeding state was not a significant predictor of blood glucose (F(1, 26.23) = 0.03, *P* = 0.867). The exercise timing × timepoint interaction remained significant after adjustment (F(1, 25.32) = 13.542, *P* = 0.00111), consistent with the primary model.

A further sensitivity model incorporating sex, age, BMI, season, and chronotype did not improve model fit (χ²(10) = 3.43, *P* = 0.9693), and none of these covariates were significant predictors of blood glucose (all *P* > 0.05). The exercise timing × timepoint interaction remained significant after full covariate adjustment (F(1, 24.88) = 13.48, *P* = 0.00115).

Consequently, the final model retained only exercise timing, timepoint, and their interaction.

### 3.4 Model 2: Acute Appetite

CAS was measured immediately before (pre) and after (post) each exercise session. Data from 27 participants were included in a primary analysis, yielding 106 observations across four condition cells (morning/evening × pre/post). Two participants were excluded due to missing CAS data across all conditions. Two further participants had one missing observation which was accommodated for by the mixed-effects framework. The LMM (Model 2, Figure 5) included exercise timing (morning *vs.* evening), timepoint (pre *vs.* post), and participant included as a random effect (*n* = 27).

**FIGURE 5:**
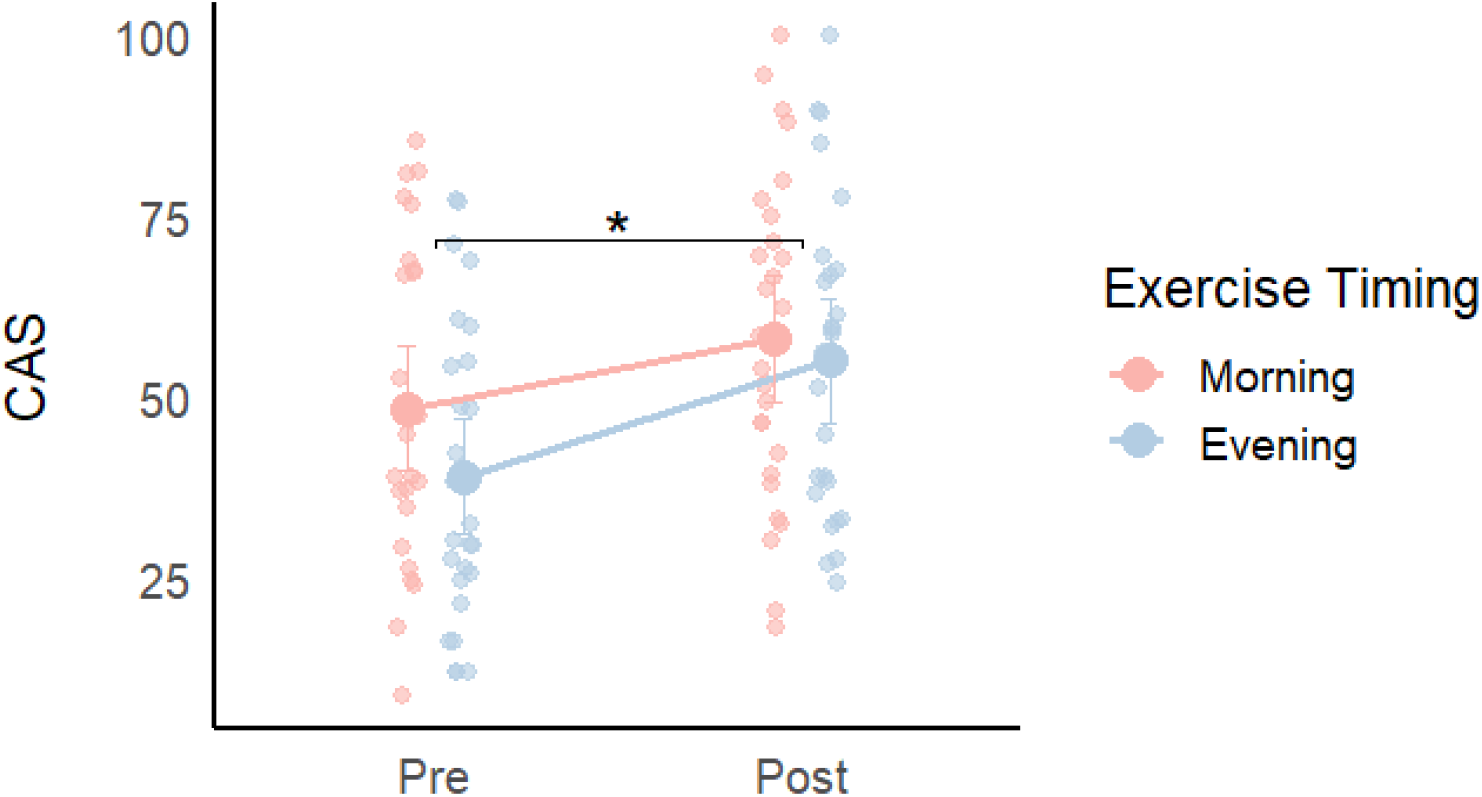
CAS pre and post exercise in the morning and evening conditions (*n* = 27). There was a significant main effect of timepoint F(1, 25.00) = 10.25, *P* = 0.00371), with CAS increasing significantly from pre- to post-exercise in both timing conditions. * *P* < 0.05.. Data presented as EMMs ± 95% CIs with individual data points.

Performing residual diagnostics indicated that there was no significant deviation from normality in the random intercept model (Shapiro–Wilk: W = 0.98, *P* = 0.0687). However, a likelihood ratio test indicated that random slopes did not produce a singular fit and significantly improved model fit (χ²(5) = 17.08, *P* = 0.00436). Subsequent diagnostics confirmed that the inclusion of slopes satisfied model assumptions (Shapiro–Wilk: W = 0.98, *P* = 0.202), and was therefore retained as the final model.

There was no significant main effect of exercise timing (F(1, 25.55) = 2.79, *P* = 0.107). The model revealed a significant main effect of timepoint F(1, 25.00) = 10.25, *P* = 0.00371), with CAS increasing from pre- to post-exercise. There was no significant exercise timing × timepoint interaction (F(1, 25.31) = 3.21, *P* = 0.0851).

A sensitivity model including a feeding state covariate (fed *vs.* fasted) did not significantly improve model fit (χ²(1) = 0.90, *P* = 0.342) and did not significantly predict CAS (F(1, 24.69) = 0.89, *P* = 0.355). The exercise timing × timepoint interaction remained non-significant after this adjustment (F(1, 25.11) = 3.26, *P* = 0.0828) which was consistent with the primary model.

A further sensitivity model including sex, age, BMI, season, and chronotype also did not improve fit (χ²(10) = 14.77, *P* = 0.141) and none of these covariates were significant predictors of CAS (all *P* > 0.05). The exercise timing × timepoint interaction remained non-significant after this covariate adjustment (F(1, 25.35) = 3.15, *P* = 0.0880).

Consequently, the final model retained only exercise timing, timepoint, and their interaction.

### 3.5 Daily Blood Glucose and Chrono-Exercise

Mean glucose, glucose SD, and CV were examined across baseline (day before exercise) and exercise day (models 3 and 4). Data from 29 participants were included in a primary analysis, yielding 112 observations across the four condition cells (morning/evening × baseline/exercise day).

Two participants contributed morning data only, resulting in a mildly unbalanced design which was supported by the mixed-effects framework. One participant had consistently low coverage across all morning observations. This participant was fully excluded from sensitivity analyses (70% coverage threshold). EMMs and 95% CIs for all outcomes across both analytical windows are presented in Table 3.

**TABLE 3.**
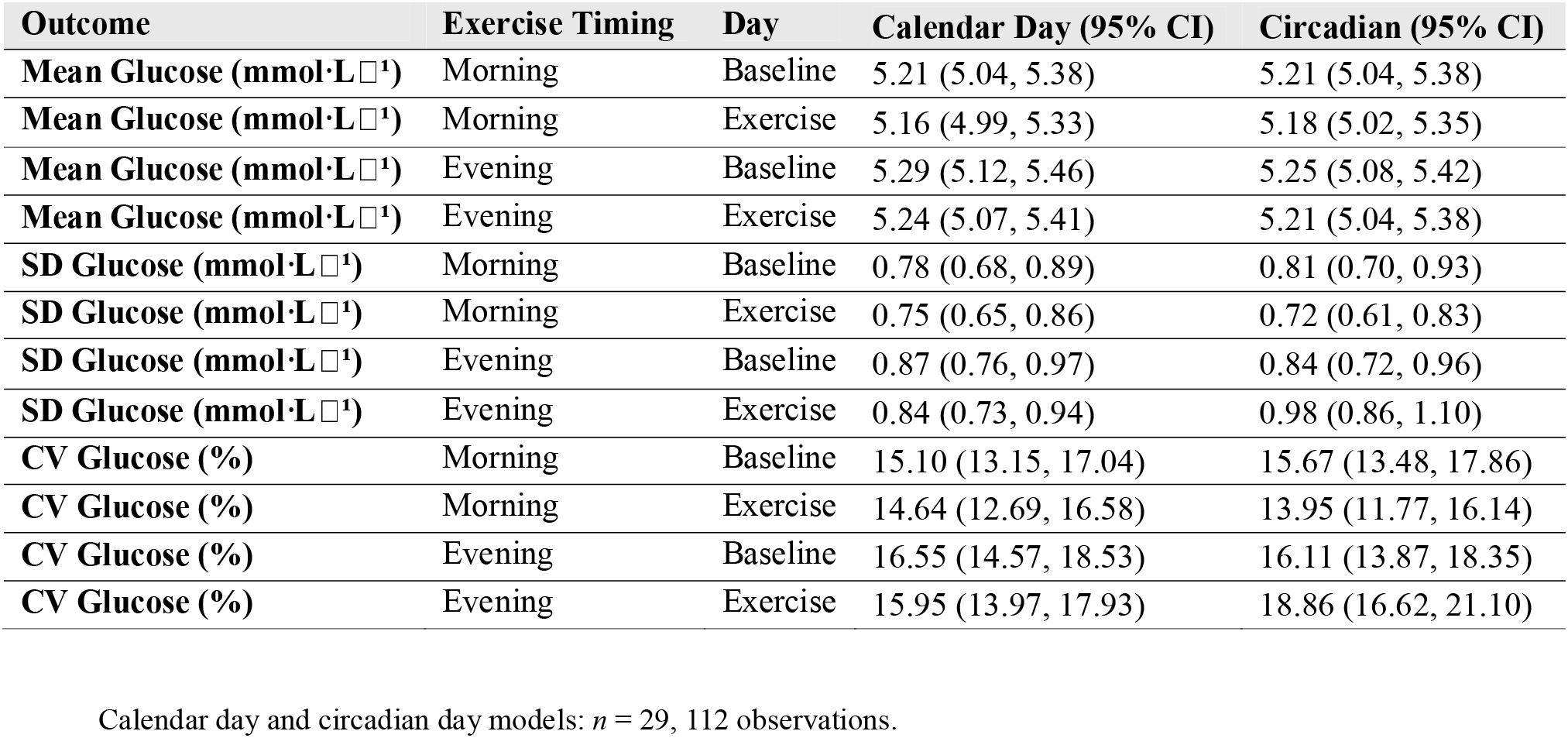
EMMs and 95% CIs for 24-Hour Glucose Outcomes Across Calendar Day and Circadian Day Analytical Windows.

#### 3.5.1 Calendar Day

Five participant-days fell below 70% coverage but were retained in the primary analysis. Mean CGM coverage was high across all conditions (morning baseline: 92.6% ± 12.9%; morning exercise day: 90.1% ± 18.7%; evening baseline: 93.1% ± 13.9%; evening exercise day: 94.4% ± 8.2%).

A likelihood ratio test indicated that random slopes did not significantly improve model fit compared to random intercepts for all three outcomes (Mean Glucose: χ²(2) = 0.032, *P* = 0.984; SD Glucose: χ²(2) = 0.074, *P* = 0.964; CV Glucose: χ²(2) = 1.037, *P* = 0.595). Random slopes also produced singular fits across all outcomes. The random intercept structure was therefore retained as the final model structure for all outcomes.

Residual diagnostics showed that there were no significant deviations from normality for SD Glucose (Shapiro–Wilk: W = 0.98, *P* = 0.107) and CV Glucose (Shapiro–Wilk: W = 0.99, *P* = 0.545). There was a significant deviation in normality observed in the mean glucose model (Shapiro–Wilk: W = 0.96, *P* = 0.00390). However, as the W statistic was close to 1.0, the deviation was considered slight. Nonetheless, bootstrap CIs (1,000 replicates) were used to validate parametric conclusions for the mean glucose model and as CIs for all fixed effects encompassed zero (day effect: −0.185, 0.094; exercise timing effect: −0.069, 0.246; day × exercise timing interaction: −0.203, 0.220), the parametric model conclusions were supported.

Sensitivity analyses applying a ≥70% CGM coverage threshold was conducted for all three outcomes under this threshold.

Figure 6 displays individual participant data and model-derived estimates.

**FIGURE 6:**
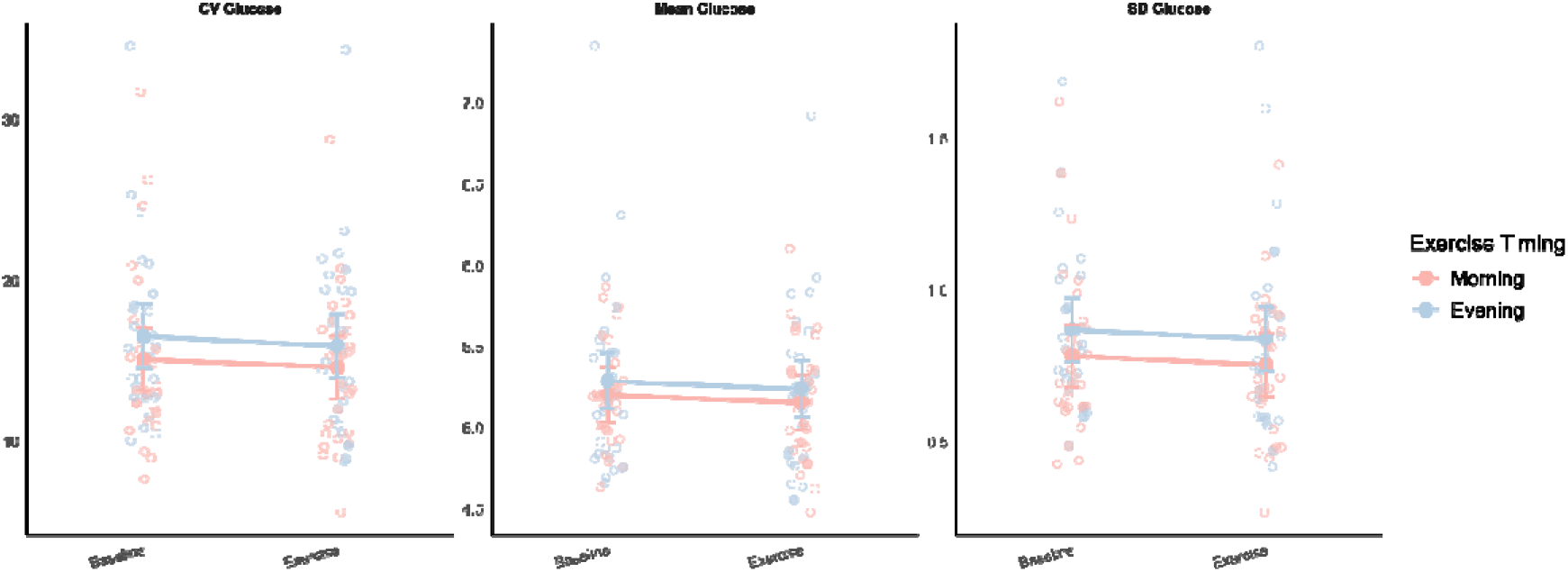
CV (%), mean glucose (mmol·L ¹), and glucose SD (mmol·L ¹), across baseline (calendar day before exercise) and exercise day in morning and evening exercise conditions (*n* = 29). There was a significant main effect of exercise timing for SD (F(1, 81.78) = 5.87, *P* = 0.0176) and CV (F(1, 81.56) = 4.85, *P* = 0.0305). Data presented as EMMs ± 95% CIs with individual data points.

##### 3.5.1.1 Model 3a: Mean Glucose

There was no significant main effect of exercise timing (F(1, 81.43) = 2.51, *P* = 0.117), day (F(1, 80.32) = 0.84, *P* = 0.361), and day × exercise timing interaction (F(1, 80.32) = 0.00, *P* = 0.993).

EMMs indicated that in the morning condition, mean glucose concentration was 5.21 mmol·L ¹ (95% CI [5.04, 5.38]) at baseline, and 5.16 mmol·L ¹ (95% CI [4.99, 5.33]) on the exercise day. In the evening condition, mean glucose was 5.29 mmol·L ¹ (95% CI [5.12, 5.46]) at baseline, and 5.24 mmol·L ¹ (95% CI [5.07, 5.41]) on the exercise day.

Applying bootstrap CIs (1,000 replicates) supported these findings as the 95% percentile CIs all encompassed zero, confirming the absence of meaningful effects on mean glucose (day; −0.183, 0.095, exercise timing; −0.072, 0.239, day × exercise timing; −0.213, 0.215).

A sensitivity model with a ≥70% CGM coverage threshold reduced the sample to 28 participants (104 observations). The results remained consistent with the primary model as there was no significant main effect of exercise timing (F(1, 73.84) = 1.56, *P* = 0.216), day (F(1, 73.33) = 0.20, *P* = 0.657), or their interaction (F(1, 73.33) = 0.26, *P* = 0.609).

A sensitivity model including feeding state as a covariate (fed *vs.* fasted) did not significantly predict mean glucose (F(1, 97.58) = 0.02, *P* = 0.880), and model adjustment did not improve fit (χ²(1) = 0.023, *P* = 0.880). The interaction remained non-significant (F(1, 79.22) = 0.00, *P* = 0.993). Therefore, feeding state was not included in the final model.

##### 3.5.1.2 Model 3b: Glucose Standard Deviation

The model revealed a significant main effect of exercise timing (F(1, 81.78) = 5.87, *P* = 0.0176) with higher overall glucose SD in the evening compared to morning (β = 0.086, SE = 0.050). There was no significant main effect of day (F(1, 80.41) = 0.74, *P* = 0.392), and day × exercise timing interaction (F(1, 80.41) = 0.00, *P* = 0.998).

EMMs indicated that in the evening exercise condition, overall glucose SD was higher (0.86 mmol·L ¹, 95% CI [0.76, 0.95]) compared to morning exercise (0.77 mmol·L ¹ 95% CI [0.68, 0.86]).

Post-hoc comparisons confirmed that there was no significant difference between baseline and exercise day within both the morning (estimate = 0.030, SE = 0.048, *P* = 0.538), and evening (estimate = 0.030, SE = 0.050, *P* = 0.550) conditions.

In the ≥70% sensitivity analysis, the exercise timing effect was non-significant (F(1, 75.11) = 3.95, *P* = 0.0506) and there were no other significant effects observed. The pattern remained unchanged, and therefore, this attenuation is most likely attributed to reduced power rather than a meaningful change.

##### 3.5.1.3 Model 3c: Glucose Coefficient of Variation

The model revealed a significant main effect of exercise timing (F(1, 81.56) = 4.85, *P* = 0.0305), with the evening condition indicating higher overall glycaemic variability relative to morning (β = 1.449, SE = 0.880).

EMMs support this finding, indicating higher overall CV in the evening condition (16.2%, 95% CI [14.5, 18.0]), compared to the morning condition (14.9%, 95% CI [13.1, 16.6]).

There was no significant main effect of day (F(1, 80.37) = 0.74, *P* = 0.394, and day × exercise timing interaction (F(1, 80.37) = 0.01, *P* = 0.911).

Post-hoc comparisons showed that there was no significant difference between baseline and exercise day within both the morning (estimate = 0.461, SE = 0.858, *P* = 0.593), and evening (estimate = 0.599, SE = 0.890, *P* = 0.503) conditions.

As with SD, the ≥70% sensitivity analysis attenuated the exercise timing effect to non-significance (F(1, 74.86) = 3.19, *P* = 0.0782) and no other significant effects were found, likely due to reduced power.

#### 3.5.2 Circadian Day

Ten participant-days fell below 70% CGM coverage but were retained in the primary analysis. Mean CGM coverage was high across baseline windows in both timing conditions (morning baseline: 93.8% ± 12.9%; evening baseline: 95.3% ± 9.6%). Coverage was more variable on exercise days, reflecting the anchoring of windows to exercise onset (morning exercise day: 82.9% ± 27.0%; evening exercise day: 84.3% ± 25.5%).

A likelihood ratio test indicated that random slopes did not significantly improve model fit compared with random intercepts for all three outcomes (Mean Glucose: χ²(2) = 0.062, *P* = 0.969; SD Glucose: χ²(2) = 0.0001, *P* = 1.000; CV Glucose: χ²(2) = 0.065, *P* = 0.968).

Random slopes also produced singular fits across all outcomes. The random intercept structure was retained as the final model structure for all outcomes.

Residual diagnostics showed significant deviation from normality across all three outcomes (Shapiro-Wilk: Mean Glucose W = 0.90, *P* < 0.001; SD Glucose W = 0.83, *P* < 0.001; CV Glucose W = 0.86, P < 0.001). Subsequently, Levene’s tests indicated that there was no significant violation of homoscedasticity for mean glucose (F(3, 108) = 1.20, P = 0.314), SD glucose (F(3, 108) = 1.21, P = 0.310), or CV glucose (F(3, 108) = 1.22, P = 0.307).

Bootstrap CIs (1,000 replicates) were used to validate parametric conclusions for the mean glucose model, and as CIs for all fixed effects encompassed zero (day effect: −0.153, 0.100; exercise timing effect: −0.114, 0.204; day × exercise timing interaction: −0.250, 0.187), the parametric model conclusions were supported.

A sensitivity analysis applying a ≥70% CGM coverage threshold was conducted for the primary outcome (mean glucose) only. Application of this threshold to secondary outcomes (SD, CV) was not pursued due to substantial data loss (10 participants).

Figure 7 displays individual participant data and model-derived estimates.

**FIGURE 7:**
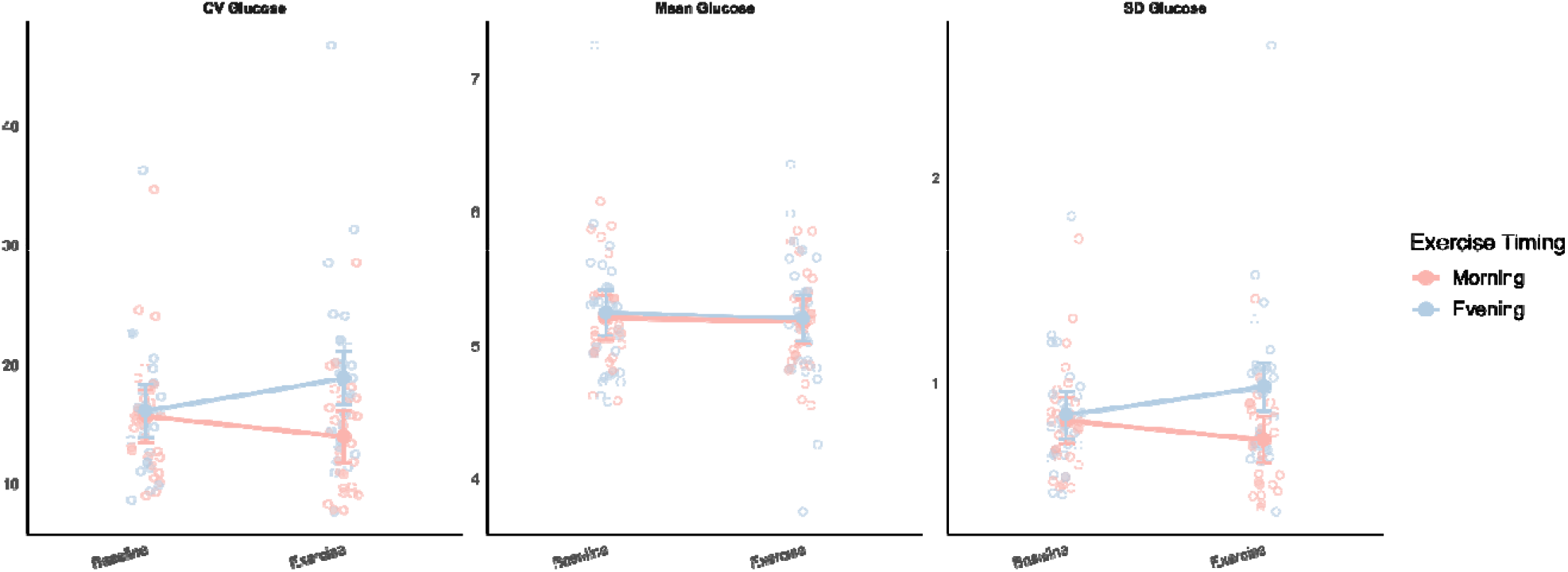
CV Glucose (%), Mean Glucose (mmol·L ¹) and SD Glucose SD (mmol·L ¹) across baseline (24 hours preceding exercise onset) and exercise day in morning and evening exercise conditions (*n* = 29). There was a significant main effect of exercise timing for SD (F(1, 82.35) = 10.71, *P* = 0.00156) and CV (F(1, 82.28) = 10.49, *P* = 0.00173). There were also significant day × exercise timing interactions for SD (F(1, 80.62) = 7.08, *P* = 0.00939) and CV (F(1, 80.62) = 7.48, *P* = 0.00766). Data presented as EMMs ± 95% CIs with individual data points.

##### 3.5.2.1 Model 4a: Mean glucose

There was no significant main effect of exercise timing (F(1, 81.44) = 0.31, *P* = 0.582), day (F(1, 80.26) = 0.43, *P* = 0.5141), and day × exercise timing interaction (F(1, 80.26) = 0.02, *P* = 0.880).

EMMs indicated that in the morning condition, the mean blood glucose concentration was 5.21 mmol·L ¹ (95% CI [5.04, 5.38]) at baseline, and 5.18 mmol·L ¹ (95% CI [5.02, 5.35]) on the exercise day. In the evening condition, mean glucose was 5.25 mmol·L ¹ (95% CI [5.08, 5.42]) at baseline, and 5.21 mmol·L ¹ (95% CI [5.04, 5.38]) on the exercise day.

Applying bootstrap CIs (1,000 replicates) supported these findings as the 95% percentile CIs all encompassed zero, confirming the absence of meaningful effects on mean glucose (day; −0.153, 0.100, exercise timing; −0.114, 0.204, day × exercise timing; −0.250, 0.187).

A sensitivity model with a ≥70% CGM coverage threshold reduced the sample to 28 participants (95 observations). The results remained consistent with the primary model as there was no significant main effect of day (F(1, 64.94) = 0.33, *P* = 0.569), exercise timing (F(1, 65.14) = 0.48, *P* = 0.493), or their interaction (F(1, 64.56) = 0.000, *P* = 0.988).

A sensitivity model including a feeding state as a covariate (fed *vs.* fasted), did not significantly predict mean glucose (F(1, 98.48) = 0.007, *P* = 0.936), and model adjustment did not improve fit (χ²(1) = 0.0073, *P* = 0.932). The interaction remained non-significant (F(1, 79.02) = 0.02, *P* = 0.880). Therefore, feeding state was not included in the final model.

##### 3.5.2.2 Model 4b: Glucose Standard Deviation

The model revealed a significant main effect of exercise timing (F(1, 82.35) = 10.71, *P* = 0.00156), with higher overall glucose SD in the evening compared to morning (β = 0.028, SE = 0.062). There was no significant main effect of day (F(1, 80.62) = 0.24, *P* = 0.622).

There was a significant day × exercise timing interaction (F(1, 80.62) = 7.08, *P* = 0.00939), indicating that the change in SD from baseline to exercise day was significantly more positive in the evening, compared to the morning (β = 0.231, SE = 0.087).

EMMs indicated that in the evening condition, there was an increase in SD from baseline (0.84 mmol·L ¹, 95% CI [0.72, 0.96]), to exercise day (0.98 mmol·L ¹, 95% CI [0.86, 1.10]). In the morning condition, SD reduced from baseline (0.81 mmol·L ¹, 95% CI [0.70, 0.93]), to exercise day (0.72 mmol·L ¹, 95% CI [0.61, 0.83]).

Post-hoc comparisons revealed that there was no significant difference between baseline and exercise day in the morning condition (estimate = 0.094, SE = 0.060, *P* = 0.123). In the evening condition, there was a significant increase in glucose SD from baseline to exercise day (estimate = −0.137, SE = 0.063, *P* = 0.0312) which supports the direction of the interaction.

##### 3.5.2.3 Model 4c: Glucose Coefficient of Variation

The model revealed a significant main effect of exercise timing (F(1, 82.28) = 10.49, *P* = 0.00173), with overall higher CV in the evening, compared to the morning. There was no significant main effect of day (F(1, 80.62) = 0.40, *P* = 0.530).

The model revealed a significant day × exercise timing interaction (F(1, 80.62) = 7.48, *P* = 0.00766). The change in CV from baseline to exercise day was significantly more positive in the evening, compared to the morning condition (β = 4.467, SE = 1.633). EMMs supported this finding showing that evening exercise was associated with an increase in CV from baseline (16.11%, 95% CI [13.87, 18.35]) to exercise day (18.86%, 95% CI [16.62, 21.10]).

On the other hand, morning exercise was associated with a reduction in CV from baseline (15.67%, 95% CI [13.48, 17.86]) to exercise day (13.95%, 95% CI [11.77, 16.14]).

Post-hoc comparisons showed that there was no significant difference between baseline and exercise day within the morning condition (estimate = 1.72, SE = 1.13, *P* = 0.1337). There was a significant increase in CV from baseline to exercise day within the evening condition (estimate = −2.75, SE = 1.17, *P* = 0.0218) which is consistent with the direction of the interaction.

## 4 Discussion

This study examined whether the timing of moderate-intensity exercise influences acute and daily glycaemic responses and subjective appetite in healthy adults under free-living conditions. Evening exercise produced a significantly greater acute reduction in blood glucose compared with morning exercise, but this did not translate into consistent differences in overall daily mean glycaemic exposure. Exercise timing was associated with differences in glycaemic variability, with the circadian-aligned analysis revealing significant day × exercise timing interactions for both SD and CV. Appetite increased following exercise regardless of timing, with no significant difference between conditions. Together, these findings suggest that exercise timing interacts with short-term glycaemic regulation and intra-day glucose dynamics in healthy adults, but its influence on appetite outcomes and overall daily mean glucose exposure appears limited in those with intact metabolic regulation.

Circadian rhythms regulate key metabolic processes, including insulin sensitivity, carbohydrate oxidation, lipid utilisation, and appetite signalling (Panda, 2016). Acute pre-exercise glucose concentrations were significantly higher in the evening than in the morning, consistent with circadian evidence that insulin sensitivity and glucose tolerance peak earlier in the day (Poggiogalle et al., 2018). Blood glucose decreased significantly from pre- to post-exercise in the evening condition, whereas no significant change was observed in the morning condition. A significant exercise timing × timepoint interaction suggests that the pattern of response differed between conditions, with a greater reduction following evening exercise compared with morning exercise. The greater reduction in the evening condition may partly reflect the higher pre-exercise glucose concentrations, with a greater capacity for reduction from a higher starting point, though the significant interaction suggests that the starting point alone does not fully account for the differential outcomes between conditions. These findings suggest that exercise interacts with circadian physiology under free-living conditions and are consistent with evidence that evening exercise may be more efficacious than morning exercise at acutely reducing blood glucose.

Daily glycaemic exposure reflects broader behavioural and physiological factors such as meal timing, sleep, habitual activity, and hormonal rhythms, and showed a more complex pattern than acute analyses. Calendar day analysis indicated no significant differences in mean glucose, SD, or CV between morning and evening conditions across baseline and exercise days for any interaction term consistent with research that found no effect of exercise timing on daily mean glucose in healthy adults (Dighriri et al., 2024). However, significant main effects of exercise timing were observed for both SD and CV, indicating that overall intra-day glycaemic variability was higher in the evening condition than the morning condition across both days. To determine whether temporal anchoring to exercise onset reveals subtler timing-related differences, a circadian-aligned analysis was also performed. Like calendar day, there was no significant difference in mean glucose, likely due to the robust homeostatic control mechanisms in healthy adults, limiting the duration and magnitude of circadian influences on glycaemia (Johnston et al., 2016; Morris et al., 2015). The main effects of exercise timing for SD and CV were strengthened and significant day × exercise timing interactions emerged for both SD and CV. Evening exercise was associated with increased glycaemic variability on the exercise day relative to baseline, whereas morning exercise was associated with a reduction. This pattern was not detectable using calendar day windows, suggesting that circadian alignment conferred greater sensitivity to these physiological changes.

CGM coverage was more variable in the circadian-aligned analysis than in the calendar day analysis, particularly for exercise days. Calendar day windows (midnight to midnight) captured a consistent daily rhythm, yielding high and stable coverage across all conditions. In contrast, coverage was more variable in circadian-aligned analyses, with high baseline coverage, compared with lower exercise day coverage. The lower coverage on exercise days compared with baseline days likely reflects exercise-related sensor disruption such as movement artefact, sweat, or temporary compression of the sensor which may in turn, reduce the frequency or reliability of readings on exercise days. Additionally, anchoring to morning exercise onset means the 24-hour window begins on average around 08:39, capturing a greater proportion of overnight hours during which sensor compression, positional changes, or app disconnection may reduce reliability. This differential coverage between conditions and between analytical approaches should be considered when interpreting the circadian-aligned findings, which may have introduced some asymmetry in the precision of glucose estimates between morning and evening conditions. As this study was restricted to baseline and exercise days, it remains unknown whether residual effects such as enhanced insulin sensitivity or altered substrate availability persisted into the post-exercise recovery period. Future studies should aim to characterise the full temporal glucose trajectory from baseline through to recovery.

CAS increased significantly from pre- to post-exercise in both the morning and the evening condition, indicating that exercise increased appetite regardless of timing. The higher mean difference in the evening condition may partly reflect the lower pre-exercise CAS than in the morning condition, differences in the time elapsed since last food intake, and diurnal variation in appetite between conditions. However, the absence of a significant exercise timing × timepoint interaction, suggests that the magnitude of appetite change did not differ between morning and evening sessions. This is broadly consistent with prior work reporting minimal effects of exercise timing on appetite responses (Maraki et al., 2005; Mode et al., 2023). The absence of a significant interaction, despite a numerically larger post-exercise increase in the evening condition (16.0 *vs.* 9.6 units), may partly reflect between-participant variability rather than a true absence of a timing effect and therefore, the marginal interaction term warrants cautious interpretation. For example, research has shown that exercise-induced hunger suppression varies according to the alignment between exercise timing and chronotype (Beaulieu et al., 2023). Although chronotype was controlled for in the present analyses, a larger sample may be required to detect potential interactions between chronotype and exercise timing. Exercise intensity may also influence appetite response as acute exercise may not always consistently elicit compensatory appetite responses (Dorling et al., 2018). Previous research has shown that acute high intensity exercise induces greater appetite suppression and reduced subsequent intake present in overweight/obese adults (Launbo et al., 2026).

The moderate-intensity protocol used here may not have been sufficient to induce appetite suppression and high intensity exercise may elicit more pronounced appetite effects, likely mediated by transient gut hormone responses, such as inducing greater ghrelin suppression (Hu et al., 2023).

Nutritional context may have obscured potential timing-related differences in appetite. Twenty-one participants completed morning exercise in a fasted state, with an average time of last intake the evening prior of 20:32, whereas all participants completed evening exercise in a fed state. The opposing influences of fasted-morning and fed-evening exercise on appetite regulation could have artificially equalised appetite scores between conditions. Feeding state may interact with exercise timing to influence postprandial glucose control while exerting little effect on appetite hormone responses or 24-hour energy intake (McIver et al., 2025), suggesting that nutritional factors may modulate glycaemic rather than appetite outcomes - consistent with the non-significant CAS interaction. In line with this, feeding state was not a significant predictor of CAS in sensitivity analyses, suggesting that the fed/fasted distinction did not statistically account for appetite differences. However, the limited variability in feeding state within conditions, with all evening participants being fed and the majority of morning participants being fasted, may have constrained the power to detect a feeding state effect. Descriptive food diary data indicated similar daily energy intake between morning and evening conditions, consistent with the null interaction finding and adding ecological validity of no systematic difference in energy intake despite *ad libitum* feeding. Nine participants did not adhere to the pre-exercise eating protocol across six morning and six evening sessions, which may have introduced some variability in nutritional state, though the sensitivity analysis suggests this did not materially influence appetite outcomes.

An important contextual factor is that this study implemented moderate-intensity exercise under unsupervised free-living conditions, chosen to reflect current physical activity guidelines recommending that adults perform weekly moderate and/or vigorous exercise (Bull et al., 2020). Many chrono-exercise studies reporting strong time-of-day effects have used vigorous-intensity or high-intensity interval training (HIIT). High-intensity exercise elicits larger metabolic and endocrine responses, including substantial glycogen depletion, elevated catecholamines, enhanced GLUT4 translocation, and prolonged improvements in insulin sensitivity (Gabriel et al., 2013; Gabriel & Zierath, 2017), which may create greater potential for circadian interactions to emerge. However, time-of-day differences have also been observed across exercise modalities including resistance training (Dighriri et al., 2025), highlighting the breadth of potential circadian influence on human physiology. Nonetheless, the moderate metabolic load may have limited the magnitude of any circadian interaction effects, particularly at the level of 24-hour integrated glucose exposure, and both glycaemic and appetite responses may be intensity-dependent. Aligning intensity with chronotype may lead to significantly greater improvements in cardiometabolic outcomes, suggesting that both timing and personalisation should be considered in exercise prescription (Tariq et al., 2026). In this study, inclusion of chronotype as a covariate did not meaningfully modify appetite or glucose responses This study captures real-world behaviour rather than laboratory-constrained physiology. Exercise intensity was verified through HR data, which confirmed that participants maintained moderate-intensity exercise in both the morning and evening conditions, with no significant difference between conditions. Participants were also able to use wearable physical activity monitors to meet a pre-defined exercise target, demonstrating the feasibility of free-living exercise prescription via wearable technology. The increasing availability of these devices provides opportunities to prescribe personalised approaches to exercise and monitor adherence in real-world settings (Terada et al., 2025). However, participant-selected outdoor routes, while ecologically valid, introduces minor variability in terrain and environmental conditions.

Whilst the physical activity monitor was also able to capture menstrual parameters, it was limited in its ability to characterise sources of potential hormonal variability across participants, which may have introduced noise in appetite and glucose outcomes. Evidence suggests that biological sex may moderate insulin sensitivity, with rhythmicity associated with metabolites modified by physical activity (MacGregor et al., 2021). Sex was included as a covariate in sensitivity analyses for models 1 and 2 and was not a significant predictor of acute blood glucose or appetite, suggesting that sex did not meaningfully predict the primary findings. However, the unequal sex distribution (18 female, 11 male) reflects recruitment constraints rather than deliberate sex-based allocation, and the study was not powered to detect sex × exercise timing interactions, limiting conclusions about sex-specific responses. Hormonal status may therefore be an important moderator of glycaemic responses to exercise timings. Future studies should recruit equal numbers of male and female participants and incorporate hormonal profiling to examine whether exercise timing effects differ between sexes and across the menstrual cycle.

CGMs allowed for dynamic analysis in a free-living setting and their use in healthy adults is increasingly recognised as feasible and informative. However, CGM systems measure interstitial rather than blood glucose, and sensor lag may increase during exercise when placed on the upper arm where muscle activity is minimal during lower-body exercise, potentially limiting the precision of acute glucose estimates. CGM accuracy may also be reduced in the normoglycaemic range where absolute glucose excursions are smaller than those observed in clinical populations, potentially limiting the sensitivity to detect subtle exercise timing effects in healthy adults. Furthermore, questions persist regarding the robustness in people not living with diabetes, including variability between company-sponsored and independent assessments of device accuracy (Oganesova et al., 2024). CGM coverage varied across participants, with some participant-days falling below 70% of expected readings, though sensitivity analyses applying a 70% coverage threshold confirmed consistent findings. Various factors such as age, sex, and exercise type may also influence CGM measurements during exercise in healthy individuals (D’Souza et al., 2023).

These limitations are intrinsic to free-living designs, but they underscore the need for future studies to integrate wearable-derived contextual data (e.g. meal timing, stress, environmental factors, sleep stage algorithms) to improve precision in modelling circadian metabolism. Despite the susceptibility of LMMs to inflated type I error rates, this study applied safeguarding methods to prevent assumption violations or misperceived random-effects structures. Residual diagnostics and applications of log transformations were used where appropriate, and results were validated by bootstrap resampling when residual normality was not achieved. Furthermore, model fit determined random-effects structures, and Satterthwaite and Kenward–Roger approximations were used to obtain more reliable estimates of degrees of freedom and standard errors. Together, these procedures reduce the likelihood of false-positive findings and increase confidence in the reported effects. The present study examined exercise timing in isolation, and future work should consider whether the modest glycaemic effects observed here are amplified when exercise timing is combined with other temporally structured interventions such as medication and nutrition timing (Peña Carrillo et al., 2024, 2026). Longitudinal interventions may reveal cumulative circadian adaptations that are not detectable from a single exercise bout. Comparing moderate and high-intensity protocols within individuals may clarify the conditions under which exercise timing most meaningfully influences glycaemic and appetite outcomes. Incorporating validated hormonal profiling alongside CGM data, and accounting for chronotype, will be important steps toward developing personalised exercise-timing frameworks grounded in circadian biology.

These findings contribute to the growing evidence that circadian biology shapes the metabolic response to exercise in healthy adults. Intrinsic skeletal muscle clocks regulate insulin sensitivity and substrate utilisation, and can be entrained by exercise as a non-photic cue (Gabriel & Zierath, 2022; Sivasengh et al., 2025). These clock-driven processes operate in healthy muscle, supporting the view that circadian exercise biology is a fundamental physiological phenomenon. Studying healthy adults therefore represents an important step in characterising the baseline circadian exercise response to inform public health guidance on exercise timing, and to provide a reference against which population-specific deviations can be understood. The ecological validity of this study, with participants maintaining free-living behaviour across the protocol, enhances the relevance of these findings to real-world exercise contexts. The circadian-aligned glycaemic variability suggests that timing may have subtle but detectable effects even in those with intact metabolic regulation, and that the restructuring of glucose dynamics around exercise is more nuanced than mean glucose measures alone can capture. Future research should examine whether these circadian-aligned variability effects are consistent across multiple exercise bouts, whether they are modified by exercise intensity or duration, and whether they extend to other metabolic outcomes and populations. In summary, these findings demonstrate that exercise timing does not influence overall daily blood glucose regulation, though distinct acute effects drive a reduction in blood glucose specific to evening exercise.

## Additional Information

### Conflicts of Interest

There are no conflicts of interest.

### Author Contributions

L.T.D. conceived the study, collected and analysed data, and drafted the manuscript. M.H., D.B., I.L., V.E., V.G., R.P., and K.B. contributed to data collection and interpretation and revised the manuscript. B.M.G. conceived and supervised the study, secured funding, and critically revised the manuscript. All authors approved the final version and agreed to be accountable for the work.

### Funding

This work was funded by the Novo Nordisk Foundation (NNF19OC0055072) and the UKRI Biotechnology and Biological Sciences Research Council (BBSRC) (BB/T00875X/1).

## Appendix

**TABLE A1.** Menstrual parameters.

| **Parameter** | **Value (*n* = 18)** |
|---|---|
| **Menstruates (*n*)** |  |
| Yes | 13 |
| No | 5 |
| **Regularity (*n*)** |  |
| Regular | 10 |
| Irregular | 3 |
| No period | 5 |
| **Cycle Length (days)** <sup>1</sup> | 33 ± 7 |
| **Period Duration (days)** <sup>2</sup> | 5 ± 1 |
| **HC (*n*)** |  |
| OCP | 2 |
| Hormonal IUD | 1 |
| Mirena Coil | 1 |

Data represent mean ± SD unless otherwise stated. Abbreviations: HC, Hormonal Contraception; OCP, Oral Contraceptive Pill; IUD, intrauterine device.
<sup>1</sup> Cycle length recorded for 12 of 13 participants who menstruate. One participant reported a cycle length of 51 days whilst self-classifying as regular. This value was retained as reported.
<sup>2</sup> Period duration calculated from the 13 participants who menstruate.

## Notes

### Competing Interest Statement

The authors have declared no competing interest.

